# Influenza C and D viral load in cattle correlates with bovine respiratory disease (BRD): Emerging role of orthomyxoviruses in the pathogenesis of BRD

**DOI:** 10.1101/2020.05.26.118166

**Authors:** Ruth H. Nissly, Noriza Zaman, Puteri Ainaa S. Ibrahim, Kaitlin McDaniel, Levina Lim, Jennifer N. Kiser, Ian Bird, Shubhada K. Chothe, Gitanjali L. Bhushan, Kurt Vandegrift, Holly J. Neibergs, Suresh V. Kuchipudi

## Abstract

Bovine respiratory disease (BRD) is the costliest disease affecting the cattle industry globally. Despite decades of research, the pathophysiology of BRD is not yet fully understood. It is widely believed that viruses predispose cattle to bacterial infection by causing direct damage to the respiratory tract and interfering with the immune system, leading to bacterial pneumonia. BRD remains a major challenge despite extensive vaccination against all major viral pathogens associated with the disease. Orthomyxoviruses (Influenza C & D viruses), have recently been found to infect cattle throughout the United States and are implicated to play a role in BRD. Here, we use the largest cohort study to date to investigate the association of influenza viruses in cattle with BRD. Cattle (n=599) from 3 locations were individually observed and scored for respiratory symptoms using the McGuirk scoring system. Deep pharyngeal and mid-nasal swabs were collected from each animal and were tested quantitatively for bovine viral diarrhea virus, bovine herpesvirus 1, bovine respiratory syncytial virus, bovine coronavirus, influenza C virus (ICV) and influenza D virus (IDV) by real-time PCR. Cattle that have higher viral loads of IDV and ICV also have greater numbers of co-infecting viruses than controls. More strikingly, in BRD-symptomatic cattle, the geometric mean of detectable IDV viral RNA was nearly 2 logs higher in co-infected animals (1.30×10^4^) than those singly infected with IDV (2.19×10^2^). This is strong evidence that viral coinfections can lead to higher replication of IDV. Our results strongly suggest that orthomyxoviruses may be significant contributors to BRD.

## 1 Introduction

Despite decades of study and attempts at control and prevention, bovine respiratory disease (BRD) remains a major cause of morbidity, mortality, and economic losses in the cattle industry worldwide. BRD is a polymicrobial infection and it accounts for approximately 70–80% of the cattle feedlot morbidity in the USA (1). Losses are estimated to be $23.60 per calf and they add up to an annual loss of over one billion dollars to the US cattle industry alone (2). BRD also results in the use of widespread therapeutics and antibiotics in feedlots, which increasingly raises public health concerns of promoting antibiotic resistance (3, 4).

The pathophysiology of BRD involves complex interactions between host, pathogen, environment, genetic and management factors. In feedlot cattle, BRD is initiated by viral infection exacerbated by stress due to travel which is typically followed by secondary infection by commensal bacteria (5). Viral infection can cause increased susceptibility to secondary bacterial infections by immunosuppression or by inflammation causing damage to the epithelium of upper airways and injuring lung parenchyma. Such damage facilitates the migration of bacterial pathogens and colonization of the lower respiratory tract. If unresolved, continued BRD advances to lower airway regions, eventually causing bronchopneumonia. Many viral pathogens have been implicated in BRD, including bovine viral diarrhea virus (BVDV), bovine herpesvirus 1 (BoHV-1), bovine respiratory syncytial virus (BRSV), bovine parainfluenza 3 (PI-3), and more recently bovine coronavirus (BCoV). In sum, more than a dozen pathogenic bacterial and viral agents have been implicated in BRD establishment and progression, prompting the development of vaccines to aid in protecting against infection with several of these organisms. However, vaccination and other prevention strategies have failed to stop the disease and alleviate these losses in production. Both bacterial and viral coinfections are common in BRD, so it can be difficult to directly implicate any one pathogen.

The latest additions to the list of BRD-associated viruses include a class of virus previously unknown to infect cattle. Orthomyxoviruses enveloped viruses with segmented negative-sense single-stranded RNA genomes, including seven genera, four of which are influenza viruses. Influenza D virus (IDV) is the most recently discovered member of the family and it is well established that cattle are the definitive host for IDV (6). IDV RNA has been found in nasal secretions from cattle with respiratory disease throughout North America (2, 6–10), Europe (11–14), and East Asia (15, 16). Although infection with IDV causes only mild respiratory disease in experimentally infected calves (17–20), a retrospective metagenomic BRD case-control study analysis of a set of 100 calves identified a positive correlation between the presence of IDV genomic RNA in nasal swabs and BRD symptoms in cattle (9). Additional retrospective case-control studies of feedlot cattle in Canada (10), the USA, and Mexico (2) have also found a higher prevalence of IDV in cattle with symptomatic BRD compared with healthy controls from similar sample sizes (n=93 and n=116). However, it remains unclear how the mild pathogenic effects of IDV infection would cause BRDC.

In 2016, a second orthomyxovirus, influenza C virus (ICV), was detected in cattle with BRD in the midwestern USA (21, 22). ICV was previously unknown to infect cattle, and it is uncertain how long ICV has existed in the cattle population. ICV has been identified in Alberta, Canada, and Texas, Oklahoma, Missouri, Colorado, Montana, Nebraska, Minnesota and Kansas, USA (10, 22). Understanding the potential role of ICV in disease in cattle is in its infancy. ICV was first identified in animals with BRD (21), suggesting that ICV may have an associative role in BRD. However, a recent retrospect case-control metagenomics study of feedlot cattle respiratory tract samples found no association between cattle with BRD and corresponding samples with reads mapping to ICV (10). Additional exploration of the role of ICV in cattle health is needed to understand the significance of this orthomyxovirus.

The potential association of these orthomyxoviruses with BRD could illuminate more about BRD pathogenesis. Previous studies of IDV/ICV in case-control cattle have included limited sample sizes and had mixed numbers of subjects from different premises. We hypothesized that exploring a larger sample-size from individual premises would provide better clarity about IDV/ICV and relationship to BRD by reducing the confounding variation contributed by differing management practices and conditions at separate premises. Since cattle serve as a natural reservoir of IDV, the mere presence of the virus may not correlate with disease. Hence, we believed that quantitative evaluation of both health status and virus copy numbers could provide a robust statistical evaluation of potential correlation of IDV or ICV with BRD. Here, probe-based real-time RT-PCRs were utilized to screen for these viruses in deep nasal swabs from cattle. This technique provides the ability to identify viral genomic RNA in a sensitive, specific and quantitative manner. Three separate case-control cohorts of approximately 200 animals each were screened to retroactively determine the prevalence of IDV and ICV in US cattle from 2011 to 2014, making this the largest case-control investigation examining the relationship between these new orthomyxoviruses and BRD.

## 2. Materials and Methods

### 2.1 Subjects/scoring/sample collection

All animal care and sample collections were approved and performed in accordance with the Institutional Animal Care and Use Committee at Washington State University (#04110). Cattle from three locations were selected for the study. One study cohort (n=200) originated from calf-raising operations in New Mexico state (23) consisting of female Holstein calves aged 23 to 76 days, sampled from August to October 2011. The other two cohorts were from beef cattle feedlots (24). One feedlot cohort (n=200) located in the state of Colorado consisted of male cattle, 98.5% Angus and 1.5% Red Angus; animals were sampled between October to November 2012. The third cohort (n=199) was from a beef cattle feedlot in Washington state with female cattle, 60.8% Angus, 14.1% Charolais, 18.1% crossbred, 4.5% Red Angus, and 2.5% Hereford; 97% of animals were sampled from January to July 2014, and 3% were sampled in October 2013. Each animal was individually observed and scored for respiratory health symptoms using the McGuirk scoring system (McGuirk, 2008), with health scores ranging from 0 to 12. Animals with McGuirk health scores ≥5 were classified as clinically affected BRD cases and those with scores ≤4 were classified as healthy controls. One deep pharyngeal and one mid-nasal swab were collected from each animal and pooled together in the viral transport medium.

### 2.2 Assay for viral agents

One aliquot of each pooled swab sample was tested for BRSV, BoHV-1, BCoV, and BVDV by the California Animal Health and Food Safety Lab System (Davis, CA) using real-time PCR or RT-PCR. Viral nucleic acids were extracted from a separate aliquot of each nasal swab using the MagMAX-96 Pathogen RNA/DNA kit following the manufacturer’s instructions. Nucleic acids were tested for PI3, IDV, and ICV using real-time RT-PCR on a 7500 Fast Real-Time PCR System (Applied Biosystems, Foster City, CA) using previously reported primer and probe sets (Supplemental Table A) (21, 25, 26) purchased from Integrated DNA Technologies (Coralville, Iowa). For PI3, 1 μL of the template was used in a 25 μL reaction using the QuantiFast Probe RT-PCR Kit (Qiagen, Inc., Valencia, CA) with 0.4 μM of each primer and 0.2 μM probe; cycling conditions were: 50 °C for 20 min, 95 C for 5 min, and 40 cycles of 95 °C for 15 s and 62 °C for 30 s.

For IDV, 8 μL of template was used in a 25 μL reaction using the AgPath-ID One-Step RT-PCR Reagents kit (Applied Biosystems) with 0.2 μM each primer and 0.06 μM probe; cycling conditions were 45 °C for 10 min, 95 °C for 10 min, and 40 cycles of 95 °C for 15 s and 64 °C for 45 s.

For ICV, 3 μL of template was used in a 20 μL reaction using the Path-ID Multiplex One-Step RT-PCR Kit (Applied Biosystems) with 0.4 μM of each primer and 0.2 μM of each probe; cycling conditions were 48 °C for 10 min, 95 °C for 10 min, and 45 cycles of 95 °C for 15 s and 60 °C for 45s.

### 2.3 Viral RNA Standards

For IDV and ICV, standard curves of transcribed RNA template corresponding to each target (IDV *PB1* gene bases 1200 to 1600 and ICV *M* gene bases 545 to 1105) were generated. RNA was transcribed from PCR products encoding the desired bases downstream of a T7 promoter region using MEGAScript T7 Transcription Kit (Invitrogen, Carlsbad, CA) followed by purification using the MEGAClear Transcription Clean-Up Kit (Invitrogen) and quantification using a NanoDrop lite instrument (Thermo Scientific, Waltham, MA). To generate standard curves, serial 10-fold dilutions of transcription product ranging from 10^−1^ to 10^8^ copies were included in triplicate in each real-time RT-PCR assay. The limit of detection for the IDV assay was 1 copy (Ct value 38.5 to 40) and for ICV it was 100 copies (Ct value 39 to 40). Three technical replicates of standards were used in all real-time RT-PCR assays, as was a negative control of nuclease-free water. Unknown samples were tested in duplicate. The coefficient of variance for all specimens (samples and standards) was Ct < 0.03.

### 2.4 Data analysis

Statistical analyses were performed using Prism version 8 (GraphPad, San Diego, CA). Mann-Whitney U tests were performed to compare clinical health scores between state cohorts, with two-tailed *p* values calculated. Viral copy numbers were determined by fitting Ct values to the standard curves generated using transcribed RNA representing the target sequence of each RT-PCR. The arithmetic mean values were calculated. To compare these lognormal-distributed viral copy data, copy numbers were log transformed and a parametric t-test using Welch’s correction was performed. In this analysis, one-tailed *p* values were calculated. Spearman correlations with two-tailed *p*-values were performed to correlate clinical scores with viral copy number; percentages were compared via a Fisher’s exact test.

## 3. Results

### 3.1 Orthomyxovirus prevalence in cattle

IDV and ICV were detected in feedlot cattle from Colorado and Washington. However, no IDV or ICV was detected in the cohort from Holstein calves from New Mexico. The prevalence of IDV was similar in both feedlot cohorts (Colorado: 5.0%, Washington: 5.5%) (Table 1). The Colorado cohort demonstrated a lower prevalence of IDV in BRD cases compared with controls (cases: 3.0%, controls: 7.1%). The Washington cohort demonstrated a similar prevalence in control and case animals (cases: 6.0%, controls: 5.1%).

**Table 1.** Significant differences (*p* < 0.05) are denoted by values with different letters).

| State | Sex | Time period | Average clinical score of cases ± SEM | Average clinical score of controls ± SEM | # tested (total/cases) | # IDV positive (total/cases) | # ICV positive (total / cases) |
| --- | --- | --- | --- | --- | --- | --- | --- |
| Colorado | Male | September – November 2012 | <sup>a</sup>7.99 ± 0.13 | <sup>a</sup>2.13 ± 0.034 | 200 / 101 | 10 / 3 | 31 / 14 |
| Washington | Female | January – July 2014 | <sup>b</sup>9.51 ± 0.13 | <sup>b</sup>2.43 ± 0.076 | 199 / 100 | 11 / 6 | 16 / 3 |
| New Mexico | Female | August – October 2011 | <sup>a</sup>8.15 ± 0.15 | <sup>a</sup>1.79 ± 0.095 | 200 / 100 | 0 / 0 | 0 / 0 |

Notably, ICV was identified in more animals than was IDV (Table 1). The prevalence of ICV in the Colorado cohort overall was 15.5% and in the Washington cohort was 8.0%. ICV was detected in 14.1% of BRD cases in Colorado and 16.8% of controls (45.2% of ICV-positive animals were cases). In the Washington cohort, 3.0% of BRD cases and 13.1% of controls were ICV-positive, with only 18.8% of ICV-positive animals having high clinical BRD scores.

### 3.2 Orthomyxovirus RNA copies in BRD case or control cattle

Although the number of BRD case animals with detectable IDV RNA (n=9) was lower than BRD controls (n=12), mean Ct values (range: control 21.82 to 37.09, case 19.70 to 34.96) and corresponding log viral copy values were different between cases and controls. Viral copy numbers were 0.75 logs higher in BRD case animals (Figure 1A). The trend toward mean viral copy numbers being higher in BRD cases compared with controls approached statistical significance (*p* = 0.097) and this trend was the same in the overall result, as well as in both of the herds independently (Colorado: *p*=0.20; Washington: *p*=0.26). In addition, a significant positive correlation was found between viral copy numbers and BRD score in the combined results (*p*=0.020) and in the Colorado cohort (*p*=0.024) (Table 2).

**Table 2.** The correlation between Influenza D viral copies and the clinical health scores.

| Cohort | Spearman r | 95% CI | P value (one-tailed) |
| --- | --- | --- | --- |
| CO | 0.65 | incalculable | 0.024 (exact) |
| WA | 0.29 | -0.39 to 0.77 | 0.19 (exact) |
| Combined | 0.45 | 0.013 to 0.75 | 0.020 (approx.) |

**Figure 1:**
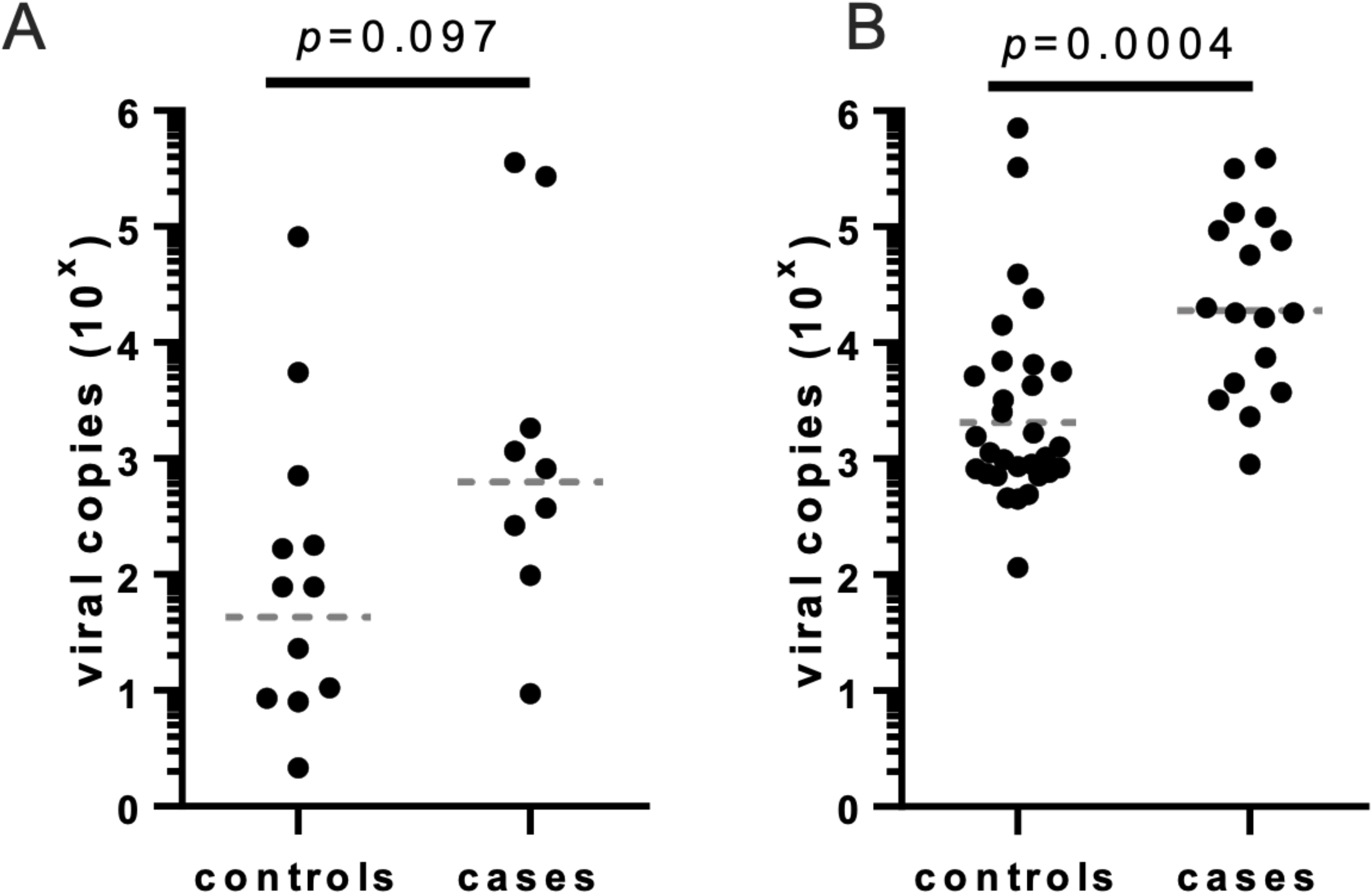
Viral RNA copy numbers from RT-PCR targeting influenza D (A) or influenza C (B) viruses were measured from nasal swabs collected from cattle demonstrating BRD symptoms (cases) or apparently healthy (controls). Each sample is represented by a circle; horizontal dotted line represents geometric mean of each population.

BRD case and control animals positive for ICV RNA both demonstrated a range of detectable Ct values (range: control 24.12 to 39.28, case 25.14 to 35.69) and copy numbers (Figure 1B). Similar to IDV, despite a higher prevalence of ICV in control cattle, the BRD case cattle mean viral copy levels were about 1 log greater, which was statistically significant in the combined cohort analysis (*p*=0.0004). As shown in Table 3, a positive correlation was found between ICV viral copy number and BRD score, which was statistically significant in the combined cohort data (Spearman r value = 0.44, *p*=0.0009) and in each cohort separately (Colorado *p*=0.022, Washington *p*=0.044), indicating that animals showing more severe clinical symptoms had more viral RNA extracted from nasal swabs.

**Table 3.** The correlation between Influenza C viral copies and clinical health score.

| **Cohort** | **Spearman r** | **95% CI** | **P value (one-tailed)** |
| --- | --- | --- | --- |
| CO | 0.36 | -0.00073 to 0.64 | 0.022 (approx.) |
| WA | 0.44 | -0.084 to 0.78 | 0.044 (exact) |
| Combined | 0.44 | 0.17 to 0.65 | 0.0009 (approx.) |

### 3.3 Association of viral presence with BRD symptoms

In addition to IDV and ICV, nasal swabs were screened for the five viruses most frequently associated with BRD namely PI3, BoHV-1, BCoV, BVDV, and BRSV. To determine if there was an association of viral presence and BRD symptoms, a Fisher’s exact test was used to evaluate the significance of odds ratio values calculated between the presence of each virus and it is a BRD case animal. The analysis showed that PI3, BoHV-1, BCoV, BRSV, and BVDV displayed odds ratios >1 (Table 4), suggesting an association of the presence of each of these viruses with BRD clinical symptoms. Of these viruses, the odds ratios of BCoV and BRSV were statistically significant by Fisher’s exact test (*p*=0.0007 and *p*=0.0008, respectively), and that of BoHV-1 approached significance (*p*=0.061). In contrast, the odds ratio for IDV was 0.74, indicating no association with BRD health status. For ICV, the odds ratio of the combined herds was 0.53, which approached statistical significance (*p*=0.062), indicating a negative correlation between the presence of ICV and the demonstration of BRD symptoms. Odds ratios of individual herds is provided in supplementary material (supplementary tables 1&2).

**Table 4.** Odds ratio of viruses with Bovine Respiratory Disease “case” status in combined results.

| **Agent** | **% positive cases (controls)** | **Odds Ratio** | **OR 95% Confidence interval (Baptista-Pike)** | **p value (Fisher's exact)** |
| --- | --- | --- | --- | --- |
| ICV | 8.5 (15.0) | 0.53 | 0.28 to 1.01 | 0.062 |
| IDV | 4.5 (6.0) | 0.74 | 0.30 to 1.73 | 0.65 |
| BVDV | 7.0 (5.0) | 1.44 | 0.64 to 3.25 | 0.40 |
| PI3 | 2.5 (1.5) | 1.69 | 0.44 to 6.46 | 0.50 |
| BCoV | 24.6 (11.6) | 2.51 | 1.46 to 4.31 | 0.0007 |
| BRSV | 6.5 (0.5) | 13.91 | 2.22 to 149.10 | 0.0008 |
| BHV-1*only in case from WA | 2.0 (0) | ∞ | 1.01 to ∞ | 0.061 |

### 3.4 Coinfections in BRD case and control animals

Each cohort investigated presented particular characteristics regarding sex, age, breed, and apparent association of viral infection with BRD clinical symptoms. While the cohort of female Holstein calves from New Mexico had a significant relationship between the presence of any virus and BRD symptoms (*p*=0.0027), animals in the feedlot cohorts, either male or female, demonstrated a significant correlation between BRD symptoms and virus presence only when 2 or more viruses were detected in the nasal swab sample (Colorado, male, *p*=0.0008; Washington, female, *p*=0.029). However, when the association of viral infection with BRD case status was assessed without counting the presence of ICV or IDV, the feedlot cattle cohorts demonstrated a significant positive correlation when at least one virus was detected (Colorado, male, *p*=0.040; Washington, female, *p*=0.0008; overall: *p*=0.0001). This finding further suggests no association between ICV or IDV and BRD clinical symptoms in these populations.

Coinfection of ICV- or IDV-positive animals with other BRD-associated viruses occurred in 22.4% of the orthomyxovirus-positive animals (Table 5). The most common virus in coinfection was BCoV (10 of the 15 coinfections). BVDV was found in coinfection in 6 of the coinfections. In the single case of coinfection with ICV and IDV, levels of both viruses were near the lower limit of detection (data not shown). ICV-positive BRD cases demonstrated a higher proportion of animals with viral coinfection compared with ICV-positive control animals (*p*=0.0016 by Fisher’s exact test), and a similar trend was observed in IDV-positive animals. In BRD case cattle, the geometric mean of detectable IDV viral RNA levels was nearly 2 logs higher in co-infected animals (1.30×10^4^) than those only infected with IDV (2.19×10^2^). However, ICV levels between these groups were similar to one another.

**Table 5.**
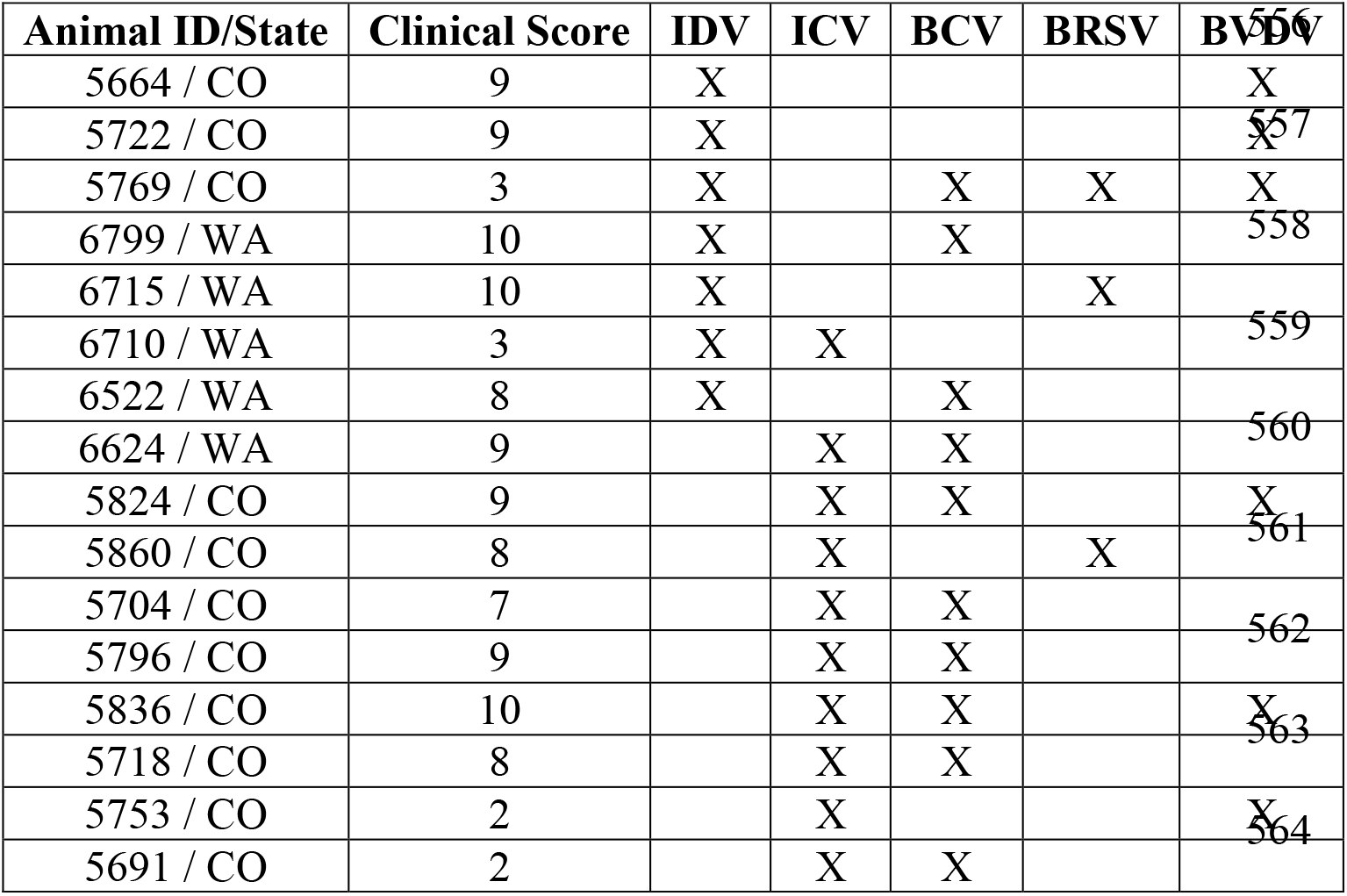
Coinfections of Influenza C and/or Influenza D positive cattle with other viruses. No animals were co-infected with with BHV-1 of P13.

## 4. Discussion

The disease of the respiratory tract in cattle is a leading cause of morbidity and mortality in the cattle industry in the US and around the world. Previous efforts to better understand the factors contributing to BRD have included genomic SNP associations, sequencing efforts, bacterial and viral associations, experimental infections with associated pathogens, as well as management interventions. Here, we discuss how the data arising from the largest case-control investigation examining the relationships among IDV, ICV, and BRD both support and dissent from this existing literature. Limitations and strengths of the different approaches are noted and directions for future research efforts are suggested.

Influenza D virus is relatively new, and it represents the first influenza virus to be closely associated with cattle (6). Indeed, cattle appear to be the reservoir of this virus, and it is known to have been present in North America since 2003 (27). ICV, which is primarily associated with disease in swine and humans, was also recently identified in North American cattle with and without BRD symptoms in the years 2015 to 2018 (10, 21). The results of this previous work provide reasonable evidence that coinfections do influence the observed highly variable clinical outcomes of BRD. More specifically, this literature provides strong evidence of higher morbidity and mortality in the event of mixed viral and bacterial infections (28). Cattle experimentally coinfected with multiple BRD-associated viruses consistently demonstrate more severe respiratory disease and prolonged viral shedding (29–31). BRD cattle coinfected with IDV and other viruses have been reported ICV, and one animal shedding IDV, ICV, BRSV, and BCoV (21). Ng *et al*. reported coinfection in six of seven IDV-positive cattle. All six coinfections were with a virus that was significantly associated with BRD in their analysis (9). In the cohorts utilized in the current study, the coinfection of IDV or ICV with other BRD-associated viruses was more frequent in symptomatic (12 out of 26 IDV/ICV-positive) versus asymptomatic (4 out of 42 IDV/ICV-positive) animals. The most common viruses found to be in coinfections with IDV were BCoV and BVDV (each in three of the six coinfections). ICV positive animals had coinfections with BCoV in seven of the nine cases and BVDV in three of nine individuals. These results strongly suggest that these orthomyxoviruses may be significant contributors to BRD by facilitating coinfections with other bovine pathogens. More strikingly, in BRD-symptomatic cattle, the geometric mean of detectable IDV viral RNA levels were nearly 2 logs higher in co-infected animals (1.30×10^4^) than those only infected with IDV (2.19×10^2^). This is strong evidence that coinfection with other viruses can lead to higher replication of IDV.

Previous studies have examined potential correlations between IDV presence and BRD symptoms and have found positive correlations with varying statistical significance. Ng *et al.* examined Californian Holstein calves (July 2011-January 2012) ages 27-60 days (9). Here, the RT-PCR method used did not include a quantitation control, so the limit of detection was unknown. Samples were reported as positive with Ct values as high as 40.39 (range 27.84 – 40.39, median 33.85). A high odds ratio based on RT-PCR results from 50 subjects of each health status was found to suggest a correlation between IDV infection and BRD symptoms, with 8 of 50 (16%) symptomatic cattle showing IDV and 0 of 50 asymptomatic animals showing ICV. Direct comparisons between the Ng *et al*. study and the current study are problematic since pathogen profiles of these 2 cohorts were different (23). Indeed, more than half of the 100 BRD case calves and one-third of control calves assayed in the current work had detectable BCoV (data not shown), but no BCoV RNA was detected in the viral metagenomic screening of the California calf cohort (9). Differences in the cohort makeup, as well as the herd management practices from these two populations may have contributed to the lack of detection of IDV in the current calf cohort. These differing results reflect the multifactorial nature of BRD.

Previous research often assayed pooled samples from different sites with variable management practices. Zhang *et al.* showed there is a wide range of BRD-associated virus prevalence associated with multiple feedlot locations (10). The current study is robust to this problem. Indeed, the analysis of large sample sizes (n=100) from two specific sights led to the same conclusions regarding the IDV and ICV correlations with BRD symptoms.

Mitra *et al.* used both metagenomics and real-time RT-PCR to detect IDV from steer at feedlots in the US state of Kansas (4 lots, total 40 animals) and multiple Mexican states (6 lots, total 53 animals) in 2015. Cattle typically enter feedlots between 4 to 6 months of age in North America. IDV was detected in one asymptomatic animal and 8 (29.6%) BRD case animals from Mexican feedlots. Using an exclusive metagenomics approach, Zhang *et al.* found a similar result of a significantly higher prevalence of IDV in samples from 58 case steer compared with 58 control steer collected from multiple feedlots in Alberta, Canada, between November 2015 and January 2016. However, in contrast to these findings supporting correlations between BRD and IDV, in the Kansas feedlots, the opposite was observed: Mitra *et al.* found IDV in only 2 asymptomatic cattle. This result is consistent with the current findings based on samples from nearly 400 individual animals from IDV-positive populations. Here, among nearly 400 individual animals from 2 different US feedlots, there was no significant evidence for higher IDV prevalence in BRD symptomatic cattle when compared with BRD asymptomatic cattle. These different correlation results amongst studies may be related to the vastly different scoring systems used to classify animals as case or control. The location (country), management practices, sampling year, age, season, and sample sizes used may all contribute to the between study correlational differences observed.

To date, only one other case-control study investigating both IDV and ICV in cattle has been reported (10). This metagenomics study, described above, found no ICV in BRD case cattle but identified several control animals with ICV. Their results agree with our finding of ICV in healthy cattle, and a higher prevalence in control animals than in cases.

Although more prevalent in healthy cattle, in the current study ICV RNA levels were significantly lower in BRD asymptomatic cattle, and IDV RNA levels showed a trend in this direction, as well. This result is in agreement with previous observations of IDV RNA levels in sick and healthy cattle (2, 32). The relative abundance of virus in each health status was not reported for ICV in the only other cattle ICV case-control report (10). Taken together, these results suggest that the level of orthomyxovirus infection may be of more relevance to BRD clinical symptoms rather than the simple presence or absence of viral RNA. It is also possible that IDV- and ICV-positive BRD control animals represent cattle with subclinical BRD. This prospect is consistent with the identification of IDV in a pig with subclinical infection in the US (33) and the observation that IDV causes only mild upper respiratory infection under experimental conditions (6, 17–20). Subclinical BRD has been identified in feedlot cattle following slaughter based on lung lesion presence and was associated with the presence of BoHV-1 in nasal-pharyngeal swabs (34). It is possible that IDV could also contribute to lung damage, as there was evidence of deep respiratory tract disease in naïve calves following experimental infection with IDV (18–20).

Unlike previous reports including several instances of coinfection of cattle with both ICV and IDV (10, 21), only one such case was identified in the current study. Previous reports of ICV with IDV coinfection were from samples obtained from 2014 and beyond, so this finding may be due to different viral dynamics during this current study (2011 to 2014). Coinfection of cattle with IDV and at least one other BRD-associated virus occurred in 55.6% of IDV-positive BRD case cattle in the current study. This is similar to the 72.2% identified by Flynn *et al* using an RT-PCR method of IDV identification (14).

Overall, a similar prevalence of RT-PCR-identified IDV and ICV was found in the BRD case and control cattle in both orthomyxovirus-positive herds tested in this study. It has been suggested (9, 14, 17) that the main role of IDV in respiratory disease is contributing to effects initiated by infection with other pathogens associated with respiratory disease in cattle. The results of this study provide further evidence to support this hypothesis. The results here suggest that while neither IDV nor ICV causes BRD directly, these orthomyxoviruses are better able to replicate in cattle with another viral respiratory infection and as such may contribute to illness in BRD. Increased replication of IDV or ICV may contribute to BRD through tissue damage. Following experimental infection of calves, IDV can be detected in samples of nasal swabs, tracheal swabs, and bronchoalveolar lavage fluid, as well as in tissues including nasal turbinate, trachea, bronchus, and lung (17–20). Inflammation in the trachea, as evidenced by neutrophil infiltration and mild epithelial attenuation, has been observed in multiple studies of experimental IDV infection in cattle (17, 19, 20). To a lesser degree, lesions have also been observed in lung tissue in some studies (18–20).

Besides physical tissue damage, the coinfection of the bovine respiratory tract with IDV and/or ICV and another virus may affect the viral replication efficiency of all coinfecting viruses. For example, in humans, infection by rhinovirus, the fastest-growing virus, reduces replication of the remaining viruses during a coinfection, while parainfluenza virus, the slowest-growing virus is suppressed in the presence of other viruses (35). Subsequently, the host is subject to the effects of the prevailing virus.

An interesting alternative possible mechanism for IDV or ICV orthomyxoviruses to affect BRD pathogenesis is through disruption of the bovine immune system. Viruses employ diverse tactics to subvert the host immune responses. For example, influenza A viruses (IAVs) can dysregulate innate immune antiviral responses in certain target cells, which promotes persistent IAV circulation in asymptomatic reservoir hosts (36–38). Particularly because IDV can be found in asymptomatic cattle, it is intriguing to consider if similar action is performed by this orthomyxovirus. A recent study showed that IDV infection could lead to suppression of cytokine production in cattle. In bronchioalveolar fluid collected two days after IDV infection, calves demonstrated upregulation of two negative regulators of cytokine production (SOCS1 and SOCS3), and the proinflammatory type one interferon pathway was decidedly unaffected (18). By suppressing immunity in the bovine respiratory tract, IDV may promote an environment permissive to other viral infections, thus promoting their establishment and facilitating the development of the disease. Delineation of the mechanism of orthomyxovirus infection of cattle leading to BRD should focus on characterizing the effects of IDV or ICV infection on bovine innate immunity.

Too often infections are studied in isolation when in reality there are most often multiple infections that all interact with the host immune system and this amalgamation results in the manifestation of the disease. As a consequence of this, the scientific community needs to embrace this added complexity as it will be necessary to clearly understand not only the infection process but also the manifestation of the disease. This approach should also inform the development of novel intervention methods. Studying processes like virus community assembly and identifying rules of the assembly could aid in the development of optimal and sustainable strategies for long term management of both animal health and productivity. This may also lead to more efficient and effective use of vaccines and antibiotics. The necessary factorial infection experiments needed to disentangle the relative roles of all of these infectious agents would be laborious and expensive. However, given the financial burden of BRD and subclinical BRD on nearly 100 million US cattle and millions more around the world, the economics warrant the availability of funding to accomplish this important animal health issue.

## 5. Author contributions

SVK conceived of the study and designed experiments. JNK and HLN conducted specimen collection. RHN, NZ, PASI, LL, KM, and JNK conducted experiments. RHN and JNK analyzed the data. IB, GLB, and SKC contributed to experiments and data analysis. RHN, KV and SVK wrote the manuscript.

## References

1. Hilton WM. 2014. BRD in 2014: where have we been, where are we now, and where do we want to go? Anim Health Res Rev 15:120–2.

2. Mitra N, Cernicchiaro N, Torres S, Li F, Hause BM. 2016. Metagenomic characterization of the virome associated with bovine respiratory disease in feedlot cattle identified novel viruses and suggests an etiologic role for influenza D virus. J Gen Virol 97:1771–84.

3. Portis E, Lindeman C, Johansen L, Stoltman G. 2012. A ten-year (2000-2009) study of antimicrobial susceptibility of bacteria that cause bovine respiratory disease complex--Mannheimia haemolytica, Pasteurella multocida, and Histophilus somni--in the United States and Canada. J Vet Diagn Invest 24:932–44.

4. Nickell JS, White BJ. 2010. Metaphylactic antimicrobial therapy for bovine respiratory disease in stocker and feedlot cattle. Vet Clin North Am Food Anim Pract 26:285–301.

5. Mosier D. 2014. Review of BRD pathogenesis: the old and the new. Anim Health Res Rev 15:166–8.

6. Hause BM, Collin EA, Liu R, Huang B, Sheng Z, Lu W, Wang D, Nelson EA, Li F. 2014. Characterization of a novel influenza virus in cattle and Swine: proposal for a new genus in the Orthomyxoviridae family. MBio 5:e00031–14.

7. Collin EA, Sheng Z, Lang Y, Ma W, Hause BM, Li F. 2015. Cocirculation of two distinct genetic and antigenic lineages of proposed influenza D virus in cattle. J Virol 89:1036–42.

8. Ferguson L, Eckard L, Epperson WB, Long LP, Smith D, Huston C, Genova S, Webby R, Wan XF. 2015. Influenza D virus infection in Mississippi beef cattle. Virology 486:28–34.

9. Ng TF, Kondov NO, Deng X, Van Eenennaam A, Neibergs HL, Delwart E. 2015. A metagenomics and case-control study to identify viruses associated with bovine respiratory disease. J Virol 89:5340–9.

10. Zhang M, Hill JE, Fernando C, Alexander TW, Timsit E, van der Meer F, Huang Y. 2019. Respiratory viruses identified in western Canadian beef cattle by metagenomic sequencing and their association with bovine respiratory disease. Transbound Emerg Dis 66:1379–1386.

11. Chiapponi C, Faccini S, De Mattia A, Baioni L, Barbieri I, Rosignoli C, Nigrelli A, Foni E. 2016. Detection of Influenza D Virus among swine and cattle, Italy. Emerg Infect Dis 22:352–4.

12. Dane H, Duffy C, Guelbenzu M, Hause B, Fee S, Forster F, McMenamy MJ, Lemon K. 2019. Detection of influenza D virus in bovine respiratory disease samples, U.K. Transbound Emerg Dis doi:10.1111/tbed.13273.

13. Ducatez MF, Pelletier C, Meyer G. 2015. Influenza D virus in cattle, France, 2011-2014. Emerg Infect Dis 21:368–71.

14. Flynn O, Gallagher C, Mooney J, Irvine C, Ducatez M, Hause B, McGrath G, Ryan E. 2018. Influenza D Virus in Cattle, Ireland. Emerg Infect Dis 24:389–391.

15. Zhai SL, Zhang H, Chen SN, Zhou X, Lin T, Liu R, Lv DH, Wen XH, Wei WK, Wang D, Li F. 2017. Influenza D Virus in Animal Species in Guangdong Province, Southern China. Emerg Infect Dis 23:1392–1396.

16. Murakami S, Endoh M, Kobayashi T, Takenaka-Uema A, Chambers JK, Uchida K, Nishihara M, Hause B, Horimoto T. 2016. Influenza D Virus Infection in Herd of Cattle, Japan. Emerg Infect Dis 22:1517–9.

17. Ferguson L, Olivier AK, Genova S, Epperson WB, Smith DR, Schneider L, Barton K, McCuan K, Webby RJ, Wan XF. 2016. Pathogenesis of Influenza D Virus in Cattle. J Virol 90:5636–5642.

18. Salem E, Hagglund S, Cassard H, Corre T, Naslund K, Foret C, Gauthier D, Pinard A, Delverdier M, Zohari S, Valarcher JF, Ducatez M, Meyer G. 2019. Pathogenesis, Host Innate Immune Response, and Aerosol Transmission of Influenza D Virus in Cattle. J Virol 93.

19. Zhang X, Outlaw C, Olivier AK, Woolums A, Epperson W, Wan XF. 2019. Pathogenesis of co-infections of influenza D virus and Mannheimia haemolytica in cattle. Vet Microbiol 231:246–253.

20. Hause BM, Huntimer L, Falkenberg S, Henningson J, Lechtenberg K, Halbur T. 2017. An inactivated influenza D virus vaccine partially protects cattle from respiratory disease caused by homologous challenge. Vet Microbiol 199:47–53.

21. Zhang H, Porter E, Lohman M, Lu N, Peddireddi L, Hanzlicek G, Marthaler D, Liu X, Bai J. 2018. Influenza C Virus in Cattle with Respiratory Disease, United States, 2016-2018. Emerg Infect Dis 24:1926–1929.

22. Zhang H, Porter EP, Lohman M, Lu N, Peddireddi L, Hanzlicek G, Marthaler D, Liu X, Bai J. 2018. Complete Genome Sequence of an Influenza C Virus Strain Identified from a Sick Calf in the United States. Microbiol Resour Announc 7.

23. Neibergs HL, Seabury CM, Wojtowicz AJ, Wang Z, Scraggs E, Kiser JN, Neupane M, Womack JE, Van Eenennaam A, Hagevoort GR, Lehenbauer TW, Aly S, Davis J, Taylor JF, Bovine Respiratory Disease Complex Coordinated Agricultural Project Research T. 2014. Susceptibility loci revealed for bovine respiratory disease complex in pre-weaned holstein calves. BMC Genomics 15:1164.

24. Neupane M, Kiser JN, Bovine Respiratory Disease Complex Coordinated Agricultural Project Research T, Neibergs HL. 2018. Gene set enrichment analysis of SNP data in dairy and beef cattle with bovine respiratory disease. Anim Genet 49:527–538.

25. Fulton RW, Neill JD, Saliki JT, Landis C, Burge LJ, Payton ME. 2017. Genomic and antigenic characterization of bovine parainfluenza-3 viruses in the United States including modified live virus vaccine (MLV) strains and field strains from cattle. Virus Res 235:77–81.

26. Faccini S, De Mattia A, Chiapponi C, Barbieri I, Boniotti MB, Rosignoli C, Franzini G, Moreno A, Foni E, Nigrelli AD. 2017. Development and evaluation of a new Real-Time RT-PCR assay for detection of proposed influenza D virus. J Virol Methods 243:31–34.

27. Luo J, Ferguson L, Smith DR, Woolums AR, Epperson WB, Wan XF. 2017. Serological evidence for high prevalence of Influenza D Viruses in Cattle, Nebraska, United States, 2003-2004. Virology 501:88–91.

28. Duff GC, Galyean ML. 2007. Board-invited review: recent advances in management of highly stressed, newly received feedlot cattle. J Anim Sci 85:823–40.

29. Potgieter LN, McCracken MD, Hopkins FM, Walker RD. 1984. Effect of bovine viral diarrhea virus infection on the distribution of infectious bovine rhinotracheitis virus in calves. Am J Vet Res 45:687–90.

30. Elvander M, Baule C, Persson M, Egyed L, Ballagi-Pordany A, Belak S, Alenius S. 1998. An experimental study of a concurrent primary infection with bovine respiratory syncytial virus (BRSV) and bovine viral diarrhoea virus (BVDV) in calves. Acta Vet Scand 39:251–64.

31. Brodersen BW, Kelling CL. 1999. Alteration of leukocyte populations in calves concurrently infected with bovine respiratory syncytial virus and bovine viral diarrhea virus. Viral Immunol 12:323–34.

32. Mekata H, Yamamoto M, Hamabe S, Tanaka H, Omatsu T, Mizutani T, Hause BM, Okabayashi T. 2018. Molecular epidemiological survey and phylogenetic analysis of bovine influenza D virus in Japan. Transbound Emerg Dis 65:e355–e360.

33. Thielen P, Nolting JM, Nelson SW, Mehoke TS, Howser C, Bowman AS. 2019. Complete Genome Sequence of an Influenza D Virus Strain Identified in a Pig with Subclinical Infection in the United States. Microbiol Resour Announc 8.

34. Kiser JN, Lawrence TE, Neupane M, Seabury CM, Taylor JF, Womack JE, Neibergs HL. 2017. Rapid Communication: Subclinical bovine respiratory disease - loci and pathogens associated with lung lesions in feedlot cattle. J Anim Sci 95:2726–2731.

35. Pinky L, Dobrovolny HM. 2016. Coinfections of the Respiratory Tract: Viral Competition for Resources. PLoS One 11:e0155589.

36. Kuchipudi SV, Tellabati M, Sebastian S, Londt BZ, Jansen C, Vervelde L, Brookes SM, Brown IH, Dunham SP, Chang KC. 2014. Highly pathogenic avian influenza virus infection in chickens but not ducks is associated with elevated host immune and pro-inflammatory responses. Vet Res 45:118.

37. Chang P, Kuchipudi SV, Mellits KH, Sebastian S, James J, Liu J, Shelton H, Chang KC. 2015. Early apoptosis of porcine alveolar macrophages limits avian influenza virus replication and pro-inflammatory dysregulation. Sci Rep 5:17999.

38. Ramos I, Fernandez-Sesma A. 2015. Modulating the Innate Immune Response to Influenza A Virus: Potential Therapeutic Use of Anti-Inflammatory Drugs. Front Immunol 6:361.

